# Rho and a riboswitch regulate *mntP* expression evading manganese stress and membrane toxicity

**DOI:** 10.1101/2023.11.29.566114

**Authors:** Anand Prakash, Arunima Kalita, Kanika Bhardwaj, Rajesh Kumar Mishra, Debarghya Ghose, Gursharan Kaur, Bibhusita Pani, Evgeny Nudler, Dipak Dutta

## Abstract

The trace metal ion manganese in excess is toxic. Therefore, a small subset of factors tightly maintains its cellular level, among which an efflux protein MntP is the champion. Multiple transcriptional regulators and a manganese-dependent translational riboswitch regulate the MntP expression. As riboswitches are untranslated RNAs, they are often associated with the Rho-dependent transcription termination in bacteria. Here we demonstrate that Rho efficiently terminates transcription at the *mntP* riboswitch region. The addition of manganese activates the riboswitch, thereby restoring the coupling between transcription and translation to evade Rho-dependent transcription termination partially. Deletion of the riboswitch abolishes Rho-dependent termination and renders bacteria sensitive to manganese due to overexpression of *mntP*. The high *mntP* expression is associated with reactive oxygen species (ROS) production, slow growth, and cell filamentation phenotypes. We posit that manganese-dependent transcriptional activation in the absence of Rho-dependent termination leads to the observed toxicity arising from excessive MntP expression, a membrane protein. Thus, we identified a novel regulatory role of Rho in preventing membrane protein toxicity by terminating at the riboswitch element.

## Introduction

Manganese (Mn) is an essential trace nutrient that regulates functions of handful enzymes either by serving as a cofactor or through differential metallation. Mn helps detoxify reactive oxygen species by acting as a cofactor of Mn-dependent superoxide dismutase and ribonucleotide reductase under iron starvation and oxidative stress ^1–3^. Mn also transiently replaces the mononuclear iron cofactor from some enzymes, and thereby restores their functions under oxidative stress^4,5^. However, in excess, Mn inhibits cell growth and promotes cell filamentation by interfering with iron homeostasis in *E. coli* ^6,7^. Mn homeostasis in *E. coli* is mainly governed by the MntR transcription regulator. Under Mn shock, Mn-bound MntR represses *mntH* and activates *mntP*, which encode Mn importer and exporter proteins, respectively ^6,8^. MntR also represses *mntS*, which encodes a small peptide, to inhibit the MntP-dependent export of Mn ^6,8^. It has been observed that the deletion of *mntP* makes *E. coli* cells highly sensitive to Mn ^6–8^. Several other mechanisms are also evolved to regulate the MntP levels in *E. coli*. For example, the expression of MntP is also regulated by Fur, a regulator that usually regulates iron homeostasis ^6^. The 5’-untranslated region (UTR) of the *mntP* forms a Mn-dependent riboswitch to upregulate *mntP* expression in *E. coli* ^9,10^. All these reports suggest that the MntP-mediated export of Mn is crucial to handle Mn homeostasis.

The riboswitches usually modulate the expression of downstream genes when they are active in the presence of small metabolites ^11,12^. Either they promote the formation of an intrinsic terminator inhibiting the transcription or act at the level of translation initiation, occluding the ribosome binding site ^10–14^. The *E. coli mntP* riboswitch belongs to the *yybP-ykoY* riboswitch family, which utilize Mn as a ligand ^10,14–17^. Without Mn, the 229 nucleotides long 5LJ-UTR of *mntP* maintains a “switched off” conformation of the riboswitch that makes RBS inaccessible to the ribosome-mediated translation initiation ^10^. The presence of Mn stimulates alternate “switch-on” conformation relieving RBS to interact with the ribosome freely ^10^. Our recent work has shown that the cellular alkaline pH favors the tight binding of Mn to the riboswitch to activate the latter. Mn activates an intrinsic alkalization circuit, overproducing cellular ammonia to attain this alkaline pH ^9^.

The Rho helicase, a hexameric transcription termination factor, binds and threads the RNA out through its central channel in a 5′ to 3′ direction. Once Rho recognizes a paused elongation complex (EC), it dissociates the RNA from the template DNA ^18–24^. Many auxiliary transcription factors can influence the Rho action. For example, elongation factor NusG binds both EC and Rho and stimulates termination ubiquitously ^25–27^. Rho function requires RNA sequences named Rho utilization (*rut*) sites, characterized by poorly conserved C-rich sequences with relatively little secondary structure ^23,28–30^. Since riboswitches are the sufficiently long untranslated RNA portion of transcripts, they frequently act as platforms for Rho-dependent termination. The Mg^2+^-sensing *mgtA*-riboswitch in *Salmonella enterica*, FMN-sensing *ribB* and *ribM-*riboswitches in *E. coli* and *Corynebacterium glutamicum*, respectively ^31–34^, lysine-sensing *lysC*-riboswitch, and thiamin pyrophosphate-sensing *thiB-*, *thiC-* and *thiM-* riboswitches from *E. coli* ^35^ are some examples where Rho-dependent termination is also reported. Despite thorough mechanistic studies, how the joint actions of riboswitch and a Rho-dependent terminator cater an evolutionary advantage to the bacteria to survive better is not addressed yet. The answer to whether the unhindered overexpression of the riboswitch-associated genes in the absence of Rho-dependent termination is detrimental to bacteria is missing.

In the present study, we show that the *mntP* riboswitch also serves as a substrate for the Rho-dependent transcription termination to downregulate *mntP* expression. Mn exposure activated the riboswitch, and moderately upregulated *mntP* expression by partially suppressing Rho-dependent termination, to mitigate Mn stress. The Mn shock in absence of Rho function overexpressed *mntP* at high level, causing membrane protein toxicity. This membrane stress perturbed oxidative phosphorylation evoking ROS production. As a consequence, bacteria exhibited slow growth and cell filamentation phenotypes. Thus, when riboswitch activation mitigates Mn stress, the Rho-dependent termination ensures healthy membrane biology by suppressing the uncontrolled expression of MntP membrane protein. Therefore, the first time we narrate the biological relevance of a Rho-dependent terminator at a riboswitch RNA region.

## Results

### Inhibition of Rho function causes upregulation of *mntP* expression

At first, we tested whether *mntP* gene expression is subjected to regulation by transcription termination factor Rho. To find a possible Rho-dependent transcription termination process upstream of any target gene, one can treat the growing *E. coli* cells with bicyclomycin (BCM), a specific and potent antibiotic against Rho function ^27,36^. Thus, we treated wild type (WT) *E. coli* strain with 100 µg/ml of BCM to inhibit Rho function. Using qPCR assay with primers located at *mntP* open reading frame (ORF) we found that BCM treatment upregulated the *mntP* RNA by 20-folds (Figure 1A). This observation suggests that the inactivation of Rho by BCM might block Rho-dependent termination at an upstream sequence, causing an increased transcriptional read-through in the *mntP* ORF.

**Figure 1.**
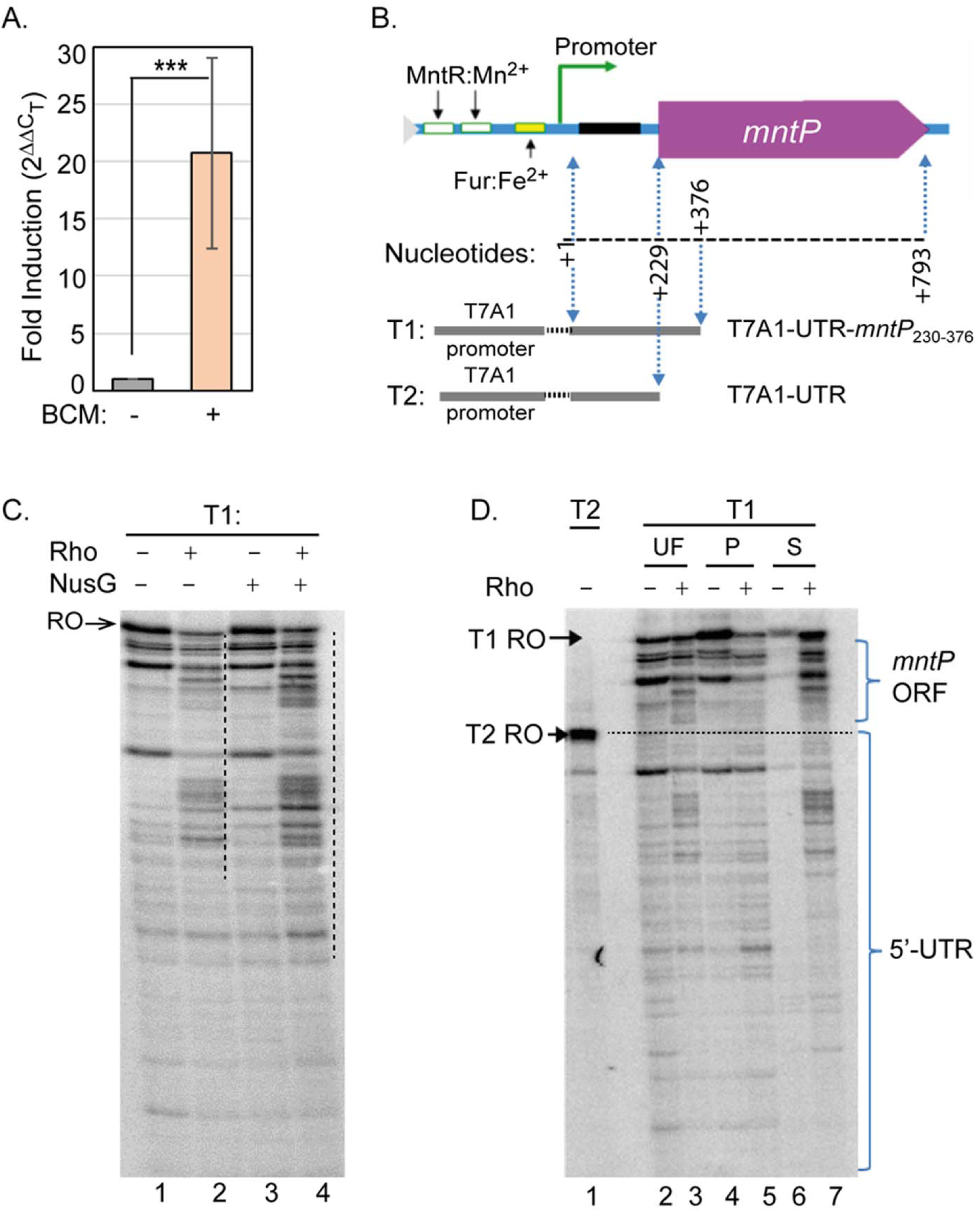
Rho-dependent termination at the *mntP* gene locus. **A.** qPCR experiment detected the upregulation of *mntP* by 100 µg/ml BCM, a Rho inhibitor. The calculated mean ± s.d. values from three independent experiments were plotted. ***P < 0.001. **B.** Schematic showing the DNA region at the 5L-UTR and *mntP* ORF that was fused with T7A1 promoter to make T1 and T2 templates for in vitro transcription. **C.** Phosphor-imaging of 6% urea PAGE showing the in vitro transcription products on the T1 template in the presence or absence of Rho, and in the presence of Rho and NusG. RO stands for Run Off products. The appearance of shorter transcripts in the presence of Rho (lane 2), and (Rho+NusG) (lane 4) were highlighted by the vertical dotted lines. **D.** Phosphor-imaging of 6% urea PAGE shows unfractionated (UF), and fractionated samples (streptavidin pellet (P) and supernatant (S) fractions) generated in the transcription assays. Comparatively shorter reaction products in the S fraction in the presence of Rho signifies that the Rho successfully terminated transcriptions. The level of RO product using a shorter T2 template (marked by the horizontal dotted line) demarcates the termination bands at the 5’-UTR and *mntP* ORF regions.

### in vitro assays confirm the Rho-dependent termination at the *mntP* gene locus

In order to check whether the BCM induced increase in *mntP* ORF transcript is a direct result of Rho action, we made an *in vitro* transcription template (T1) fusing T7A1 promoter with 5’-UTR and 147 bp of *mntP* ORF by an overlapping PCR (Figure 1B). A single round *in vitro* transcription assay was performed in the presence or absence of Rho. The transcription products were resolved by a urea-denaturing polyacrylamide gel electrophoresis (Urea-PAGE). Many prominent transcript bands shorter than the full-length read-through RNA transcripts appeared, when transcription was performed in the presence of Rho (Figure 1C, lane 2). These bands represent transcription termination products. NusG is an additional transcription factor that assists the Rho-dependent termination process ^25–27^. We found that NusG enhanced the band intensities and early appearance of the shorter transcripts in the presence of Rho (Figure 1C, lane 4).

Although the shorter transcripts are considered the putative Rho-dependent transcription termination products, they could be the mixture of the actual products of termination and transcriptional pausing of ECs (Figure 1D). To separate the paused and terminated transcripts, we performed transcription assays immobilizing biotinylated-template on streptavidin beads. The reaction products were separated into pellet and supernatant fractions. In the absence of Rho, most of the transcripts appeared in the pellet, while barely any transcripts were visible in the supernatant fraction in a urea-PAGE (Figure 1D, lanes 4 and 6). However, when Rho was present, most of the shorter transcripts appeared in the supernatant than in the pellet fraction (Figure 1D, lanes 5 and 7). The appearance of transcripts in the supernatant fraction indicates that Rho successfully terminated many transcripts before maturation. A shorter transcriptional read-through product using the T7A1-UTR template (T2) was generated as a marker (Figure 1B and 1D). The Rho-dependent transcription termination bands that were resolved below or above the position of the T2 RNA marker represent the termination that occurred in the 5’-UTR or 147 bp of *mntP* ORF, respectively (Figure 1D). This data elucidates that Rho terminates transcription at the 5’-UTR as well as at the ORF region of *mntP*.

### The activated riboswitch partially overcomes Rho-dependent termination

We then asked how Mn affects Rho dependent termination, or vice versa, at the *mntP* locus. Mn regulates both the *mntP* riboswitch and the *mntP* promoter ^6,10^. Therefore, we chose to fuse the riboswitch region with the T7A1 promoter to get reporter constructs for cleaner experimentations. We designed three reporter constructs encompassing various lengths of DNA corresponding to the 5’-UTR of *mntP* to dissect the effects of Mn and Rho in the riboswitch activation (Figure 2A). T7A1 promoter followed by the 5’-UTR and a portion of *mntP* ORF (encompasses +1 to +439 bp) was transcriptionally fused with either *lacZ* or YFP reporters. The reporter cassettes were integrated in the WT *E. coli* chromosome to get the construct 1 and 2 strains of *E. coli* (Figure 2A). Consistent with the qPCR trend (Figure 1A), BCM treatment (100 µg/ml) enhanced the β-galactosidase activity and YFP fluorescence up to about 23 and 28-folds, respectively (Figures 2B and 2C). This observation suggests that in vivo Rho-dependent termination at the *mntP* locus suppresses reporter expressions.

**Figure 2.**
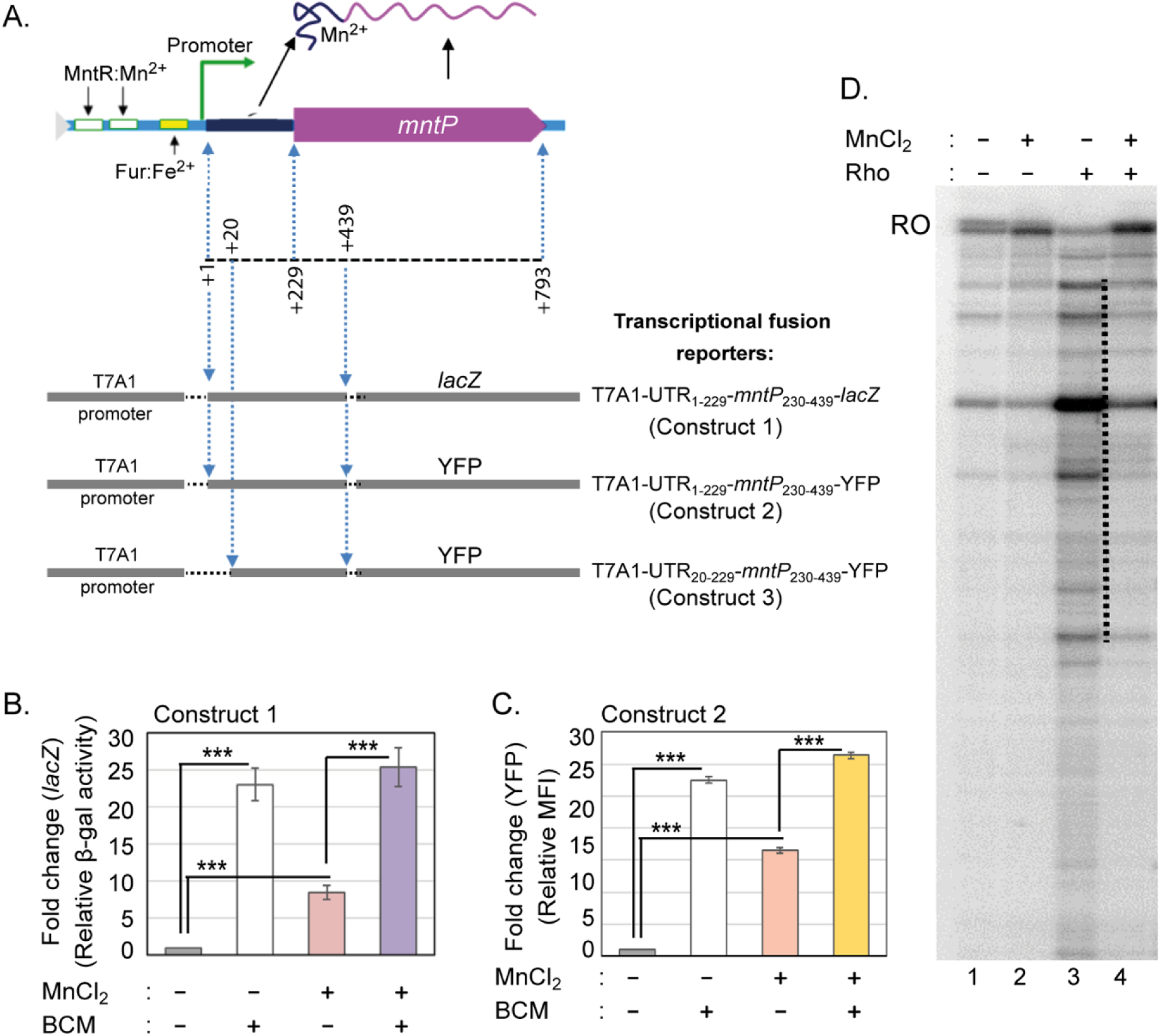
The reporter assays to demonstrate that Rho prematurely terminates *mntP* expression. **A.** The schematic shows the DNA fragments from the 5L-UTR and *mntP* ORF that have been fused with T7A1 promoter and YFP or lacZ reporters, as indicated. to make the reporter cassettes. The reporter cassettes were individually integrated with the WT *E. coli* genome to get the constructs 1, 2, and 3. **B.** The estimated β-galactosidase activity from construct 1 in the presence or absence of 100 µg/ml BCM and 8 mM Mn suggests that Rho terminates transcription prematurely in the presence or absence of the activated riboswitch. The calculated Miller unit for construct 1 was 545.5 ± 41. **C.** The estimated MFI of YFP from construct 2 exhibited similar trends as β-galactosidase activity (Panel B) to suggest that Rho terminates transcription prematurely both in the presence or absence of the activated riboswitch. The calculated mean ± s.d. values from three independent experiments were plotted. ***P < 0.001. **D.** Phosphor-imaging of in vitro transcription products on the T1 template in the presence or absence of Rho, and/or 100 µM Mn indicates that Mn inhibits Rho dependent termination. RO stands for Run Off products.

We also tested the effect of Mn on the expression profile of reporter constructs 1 and 2. The MnCl_2_ (8 mM) shock (since 8 mM exogenous Mn causes sufficient toxicity in *E. coli* WT cells (18)) enhanced both β-galactosidase activity (9-folds) and YFP fluorescence (16-folds) (Figure 2B and 2C). BCM treatment to the Mn-fed *E. coli* cells further enhanced the β-galactosidase activity (25-folds) and YFP fluorescence (30-folds) that are comparable to the levels achieved by BCM treatment alone (Figure 2B and 2C). This observation indicates that while the Mn-mediated riboswitch activation upregulates the reporter genes partially, most of the transcripts still remain under regulation by Rho dependent termination. In other words, Mn could partially inhibit Rho dependent-termination by activating *mntP* riboswitch to upregulate the reporter genes.

To address whether Mn directly could affect Rho-dependent termination at the *mntP* riboswitch, we employed in vitro transcription assay to test termination in a purified system. Upon addition of Mn, longer transcripts representing transcription run off re-appeared in reaction containing termination factor Rho (Fig 2D). This result reinforced the in vivo observation that *mntP* 5’-UTR serves as a Mn-dependent riboswitch by dialing down Rho dependent termination.

### Free Mn does not influence Rho’s function

To test whether free Mn directly affects Rho’s termination function, we designed the following experiments. From the predicted structure of *mntP* riboswitch (10), we hypothesized that the absence of +1 to +19 nucleotides of the 5’-UTR would diminish the riboswitch activation while allowing synthesis of a sufficiently long untranslated RNA for Rho-dependent termination. To test this possibility, we generated the 3^rd^ reporter strain (construct 3) where the T7A1 promoter followed by +20 to +439 bases of the *mntP* gene was fused with a YFP reporter, and integrated into the WT strain (Figure 2A). Mn at 8 mM did not affect the mean fluorescence intensity (MFI) of YFP in the construct 3 (Figure 3A). However, BCM alone, or BCM treatment to the Mn-fed construct 3 strain, increased the MFI of YFP to 15-fold (Figure 2D). This observation indicates that the free Mn that could not form complex with the truncated riboswitch, has no effect on Rho function, and BCM-mediated inactivation of Rho directly allows transcription through the 5’-UTR to express YFP reporter.

**Figure 3.**
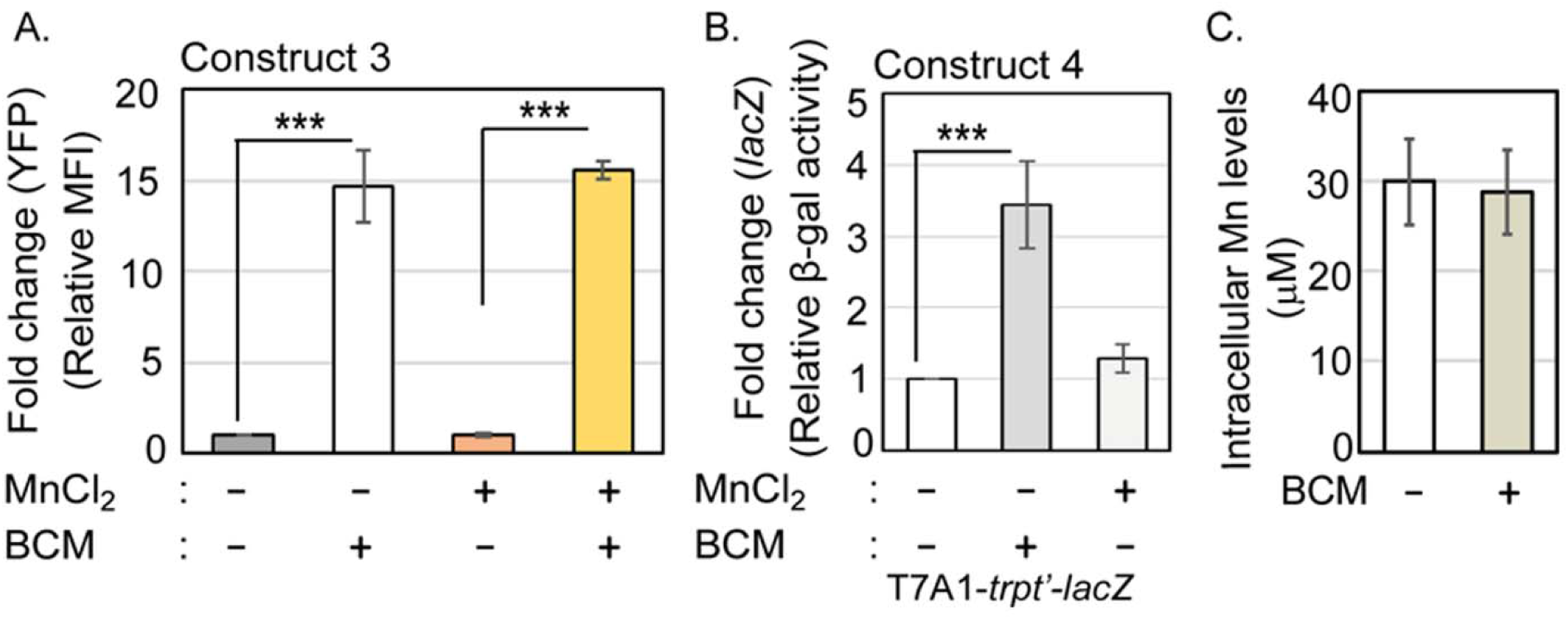
Free Mn and Rho do not affect each other’s actions. **A.** Mn (8 mM) has no effect on YFP reporter activation in construct 3, which contains a slightly truncated *mntP* riboswitch that allows Rho-dependent termination but inactivates Mn-dependent riboswitch function. However, MFI of YFP reporter in the presence of 100 µg/ml BCM indicates that Rho efficiently terminates YFP reporter expression both in the presence or absence of Mn. **B.** BCM (100 µg/ml) treatment increases the β-galactosidase activity in construct 4 containing *trp t’* terminator (a tryptophan operon Rho terminator). However, Mn treatment (8 mM) did not increase β-galactosidase activity, suggesting that Mn does not inhibit Rho-dependent termination. The calculated Miller unit for construct 4 without any treatment was 4428 ± 902. **C.** BCM (100 µg/ml)-mediated Rho inactivation does not alter cellular Mn level, suggesting that Rho termination has no impact on Mn homeostasis. The calculated mean ± s.d. values from three independent experiments were plotted. ***P < 0.001.

To further ensure that Mn does not directly affect Rho function, we made construct 4, where the T7A1 promoter was fused with *trp t’* DNA that encodes a well-known Rho-terminator site ^20,24,37^, and a *lacZ* reporter. Using this construct, we show that 8 mM Mn did not affect the Rho-dependent termination process (Figure 3B). Besides, performing ICP-MS analyses, we determined that inactivating Rho-dependent termination by BCM did not increase the cellular Mn levels (Figure 3C). All these results suggest that Mn shock and Rho-dependent termination do not directly influence each other to affect cellular physiology.

### *E. coli* is Mn-sensitive in the absence of the 5’-UTR

We wondered why two opposing of regulations were operated at the same region during gene expression, viz., Rho-dependent silencing and riboswitch-dependent activation, are evolved to mediate *mntP* gene expression simultaneously. To address this question, we deleted the riboswitch (except +1 to +20 bp region) from the WT strain to get Δ*RS*::*kan^R^*and Δ*RS* strains by the λ-red recombination system ^38^. Surprisingly, the riboswitch-deleted strains were Mn sensitive when allowed to grow on an LB-agar plate supplemented with 2 mM MnCl_2_ (Figure 4A). The degree of Mn susceptibility of the riboswitch-deleted strains appeared to be milder than the Δ*mntP* strain, a known Mn-sensitive strain of *E. coli*, which is highly sensitive under 1 mM MnCl_2_ ^7^; and more susceptible than the WT strain under the same condition (Figure 4A). Performing growth curve analyses, we find that 1 mM and 3 mM Mn caused roughly equivalent growth defects to the Δ*mntP* and the Δ*RS* strains, respectively (Supplementary Figure 1).

**Figure 4.**
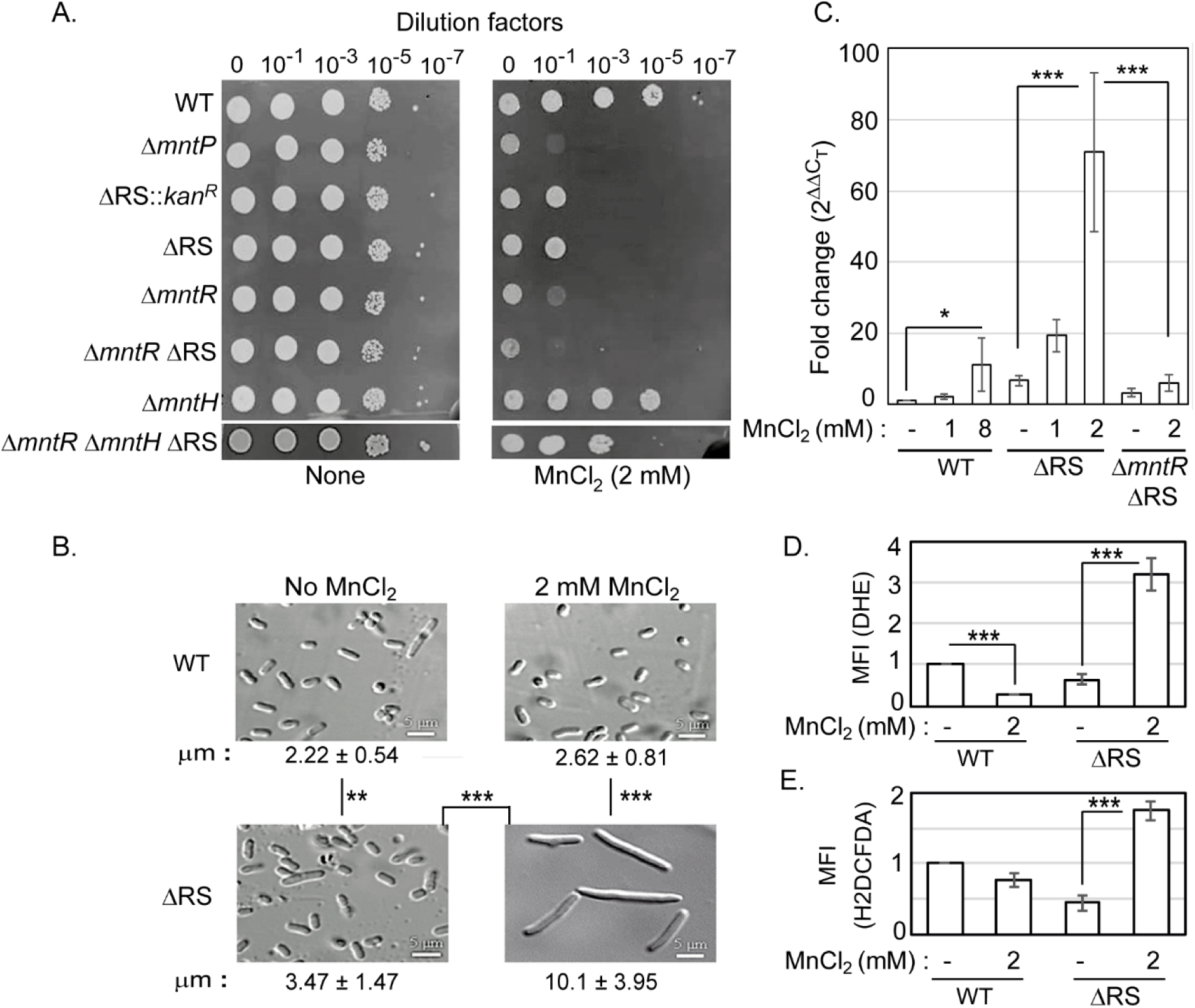
Rho-dependent termination at 5’-UTR inhibits membrane protein toxicity. **A.** The riboswitch-deleted strains (Δ*RS::kan^R^*and Δ*RS*) are Mn sensitive. The absence of MntR in the Δ*mntR* Δ*mntH* Δ*RS* strain partially helped to overcome Mn toxicity. **B.** Confocal microscopy images indicate that the Δ*RS* strain is significantly elongated than the WT strain. 2 mM Mn promotes cell filamentation of the Δ*RS* strain. The µM length of cells were calculated (mean ± s.d. values from 30-50 cells) and shown. **P < 0.01; ***P < 0.001. **C.** qPCR data show that the *mntP* gene expression in the untreated Δ*RS* strain is naturally higher than the WT strain. 1 mM and 2 mM Mn-treated Δ*RS* strain increased the *mntP* expression, which are greater than the expression in the 8 mM Mn-treated WT strain. Mn failed to induce *mntP* expression in the Δ*mntR* Δ*RS* strain. *P <0.1; **P <0.01. **D.** The ROS detecting probe DHE fluorescence (MFI) was significantly decreased in WT cells treated with Mn. However, the fluorescence was increased in the Mn-treated Δ*RS* strain. **E.** The ROS detecting probe H2DCFDA fluorescence (MFI) was slightly decreased in WT cells treated with Mn. However, the fluorescence was increased in the Mn-treated Δ*RS* strain. The calculated mean ± s.d. values from three independent experiments were plotted. ***P < 0.001.

To know whether Mn sensitivity affects cellular morphology, we performed confocal microscopy. The WT strain exhibited a typical average cell length (2.22 ± 0.54 µM). In comparison, the Δ*RS* strain showed a significant increase in average cell length (3.47 ± 1.47 µM) in LB broth (Figure 4B). Under 2 mM Mn shock, the WT strain showed a significant but minor increase in average cell length (2.62 ± 0.81 µM), while the Δ*RS* strain was filamented (10.1 ± 3.95 µM) (Figure 4B). Filamentous phenotype of ΔRS strain was further analyzed by FM-4-64 and DAPI staining to show that the nucleoids were segregated but there was no septum formation in the filamentation (Supplementary Figure 2). Interestingly, the Δ*mntP* strain has also exhibited Mn-dependent cell filamentation ^7^. In summary, both absence of *mntP* (in the Δ*mntP* strain) and absence of Rho-dependent termination/riboswitch function (in the riboswitch-deleted strains) caused stressed phenotypes under Mn shock.

### Rho-mediated silencing of *mntP* is crucial to evade membrane protein toxicity

Generally, the overexpression of inner membrane proteins is highly toxic and leads to growth defects and cell filamentation ^39–42^. Since MntP is an inner membrane protein, it was very likely that the observed growth defects and cell filamentation (Figure 4) could be the result of excessive overexpression of MntP in the Mn-stressed ΔRS strains. If so, the mechanism of such *mntP* overexpression could be originated by two simultaneous incidents, viz. absence of Rho-dependent termination and activation of MntR activator by Mn in the Mn-fed Δ*RS* strains (Figures 4A and 4B). Therefore, we generated Δ*mntR* Δ*RS* double mutant strain to nullify the MntR-mediated regulation. However, the Δ*mntR* Δ*RS* could not rescue the growth defect under Mn stress (Figure 4A), possibly because as the Δ*mntR* strain is Mn sensitive ^6^, therefore, Δ*mntR* Δ*RS* was also sensitive to Mn. Upregulation of *mntH*, a high affinity Mn importer, in the Δ*mntR* Δ*RS* strain, could facilitate Mn import to make the strain sensitive under Mn shock. Indeed, we show that a triple mutant, Δ*mntR* Δ*mntH* Δ*RS*, partially rescued the growth defects under Mn shock (Figure 4A).

We probed the transcriptional upregulation of *mntP* in *E. coli* strains by qPCR analyses. Mn at 1 mM, which is nontoxic to the WT cells (18), slightly enhanced the expression of *mntP* (2.5-folds) (Figure 4C). When treated with a toxic dose (8 mM) of Mn ^9^, the WT cells exhibited about 11-fold upregulation of *mntP* (Figure 4C). The Δ*RS* strain showed an inherently high level of *mntP* expression (7-folds) even in the absence of Mn shock (Figure 4C). Strikingly, Mn at 1 mM and 2 mM levels increased this expression to about 19 and 70-folds, respectively (Figure 4C). As expected, unlike the Δ*RS* strain, Δ*mntR* Δ*RS* strain respectively exhibited 3-fold and 6-fold upregulation of *mntP* compared to the WT strain in the absence and the presence of 2 mM MnCl_2_ (Figure 4C). The upregulation of *mntP* in the Mn-treated the Δ*RS* strain was found to be associated with oxidative stress, as observed using dihydroethidium (DHE) and 2’,7’-dichlorodihydrofluorescein diacetate (H2DCFDA), two different ROS detecting chemicals (Figure 4D and 4E). This data is consistent with a previous observation that has shown that the *mntP* overexpression from a plasmid leads to oxidative stress in *E. coli* ^43^.

To directly check whether *mntP* overexpression is toxic, we transformed the Mn-sensitive Δ*mntP* strain with an arabinose inducible pBAD-*mntP* expression vector. The multicopy leaky expression of *mntP* from the plasmid sufficiently rescued the Mn-sensitive growth phenotype of the Δ*mntP* strain (Supplementary Figure 3A). The Δ*mntP* strain harboring the pBAD empty vector did not rescue the Mn-dependent growth inhibition (Supplementary Figure 2A). However, when *mntP* expression from the pBAD-*mntP* expression vector was induced by 0.002% arabinose, the Δ*mntP* strain showed extreme growth retardation (Supplementary Figure 2A). We also tested whether, similar to the Mn-fed Δ*RS* strain (Figure 4B), *mntP* overexpression leads to cell filamentation. Indeed, the multicopy overexpression of *mntP* led to cell filamentation (Supplementary Figure 3B). Further, to check whether excessive Mn export by overexpressed MntP leads to cell filamentation phenotype, we used a nonfunctional MntP mutant ^43^ (pMntP^D118A^ transformed in the the Δ*mntP* strain) in the growth assay. The multicopy induction of MntP^D118A^ by arabinose (0.002%) did not rescue the growth defect (Supplementary Figure 3C), suggesting that MntP overexpression but not its function is toxic to the cells.

We attempted to check the levels of MntP induced by arabinose (0.002% and 0.02%) in the WT strain harboring the pBAD-*mntP* vector. We also transformed the pET-*mntP* vector, which was another construct made to produce MntP with C-terminal his_6_ tag, in the *E. coli* BL21 (DE3), C43(DE3) and Lemo (DE3) strains to check MntP-6X-his overexpression by 0.5 mM IPTG. Primarily, C43 (DE3), and Lemo (DE3) strains are known to suppress membrane protein overexpression related toxicity ^44,45^. However, the induction of MntP was not detected in the SDS-PAGE (Supplementary Figure 3D and 3E). In the western blot experiment, the overexpressed MntP-6X-His from the pET-*mntP* vector also remained undetected against a polyclonal anti-6X-His antibody (data not shown). These observations indicate that the overexpressed MntP could be highly toxic even below detection limit (BDL). This overexpression of MntP at BDL is also apparent in a previous study that could barely able to detect the MntP overexpression by performing a western blot experiment even after using a C-terminal 3X-FLAG epitopes ^6^. Taking all observations in account, we hypothesize that the Rho function ensures a low level of *mntP* expression to evade membrane protein toxicity under Mn stress.

## Discussion

Our current study demonstrates that Rho is pivotal in terminating the transcription of *mntP* at the riboswitch. In the presence of Mn stress, the active Mn^2+^:*mntP*-riboswitch is formed that switches on the translation ^10^, thereby partially evading Rho-dependent transcription termination and allowing MntP expression (Figure 2B and 2C). Thus, efflux of Mn by overexpressed MntP mitigates manganese stress.

A tight coupling between transcription and translation processes, in vivo, usually evades the Rho-mediated termination at the ORF ^46,47^. In the absence of Mn, riboswitch’s “switched off” conformation masks the SD site ^10^, and thereby former would uncouple the transcription and translation processes. Thus, Rho-dependent termination suppresses roughly 24/25^th^ parts of the reporter gene expression, while only 1/25^th^ of parts could express (Figure 2B and 2C). In the presence of Mn shock, the alternative “switch on” conformation of riboswitch would evade Rho-dependent transcription termination in two different ways: one is by directly inhibiting Rho-binding to the RNA, and the other is by unmasking the SD, thereby facilitating transcription and translation coupling. Thus, under Mn stress, a modest increase (about 8/25^th^ parts) in β-galactosidase activity was observed (Figure 2B and 2C).

While most riboswitches from *Bacillus subtilis* affect intrinsic termination ^11,48^, the *E. coli* riboswitches work primarily by modulating translation initiation ^49^. Consistently, the *mntP* riboswitch of *E. coli* work at the level of translation initiation ^10^. Interestingly, majority of the *E. coli* riboswitches employ Rho-dependent transcription termination ^31,35^. Rho is abundant and essential in *E. coli*; therefore, the scope of Rho-dependent termination is widespread in *E. coli* ^23,27,50^. On the other hand, Rho is scarce and dispensable in *B. subtilis*; therefore, most of the terminators are intrinsic in *B. subtilis* ^51,52^. Thus, generalizing others’ ^31,35^ and our current observations, it appears that the active riboswitches in *E. coli* defy the Rho-dependent termination by coupling transcription and translation processes, and thereby allow gene expression. While in *B. subtilis*, as the Rho-dependent termination is not a norm, the riboswitches modulate intrinsic termination function for gene expression. Whether this generalization has a broader scope for Gram-negative and Gram-positive bacteria needs to be deciphered.

Despite the riboswitch suppresses Rho-dependent termination, the bulk of total transcription (about 17/25^th^ parts) was still prematurely terminated by Rho (Figure 2B and 2C). We argued that most of the time, Rho quickly gets access to the nascent free RNA as it migrates with the EC ^24^ before Mn could form a complex with RNA to “switch on” the riboswitch under in vivo conditions. This argument explains how Rho-dependent termination at the *mntP*-riboswitch region ensures a substantially low level of MntP expression both in the presence or absence of Mn stress. Why does Rho warrant low level expression of MntP? To answer this question, we demonstrate that when Rho-dependent termination at the riboswitch RNA was defied using an Δ*RS* strain, an inherently high level of *mntP* expression was detected (Figure 4C). Mn stress increased this expression further (82-fold) by activating the MntR regulator (Figure 4C), eliciting membrane protein toxicity. Therefore, we propose that the Rho-dependent transcription termination acts as a guardian of the cell against membrane protein toxicity by silencing MntP expression under Mn stress, as shown in the schematic (Figure 5).

**Figure 5.**
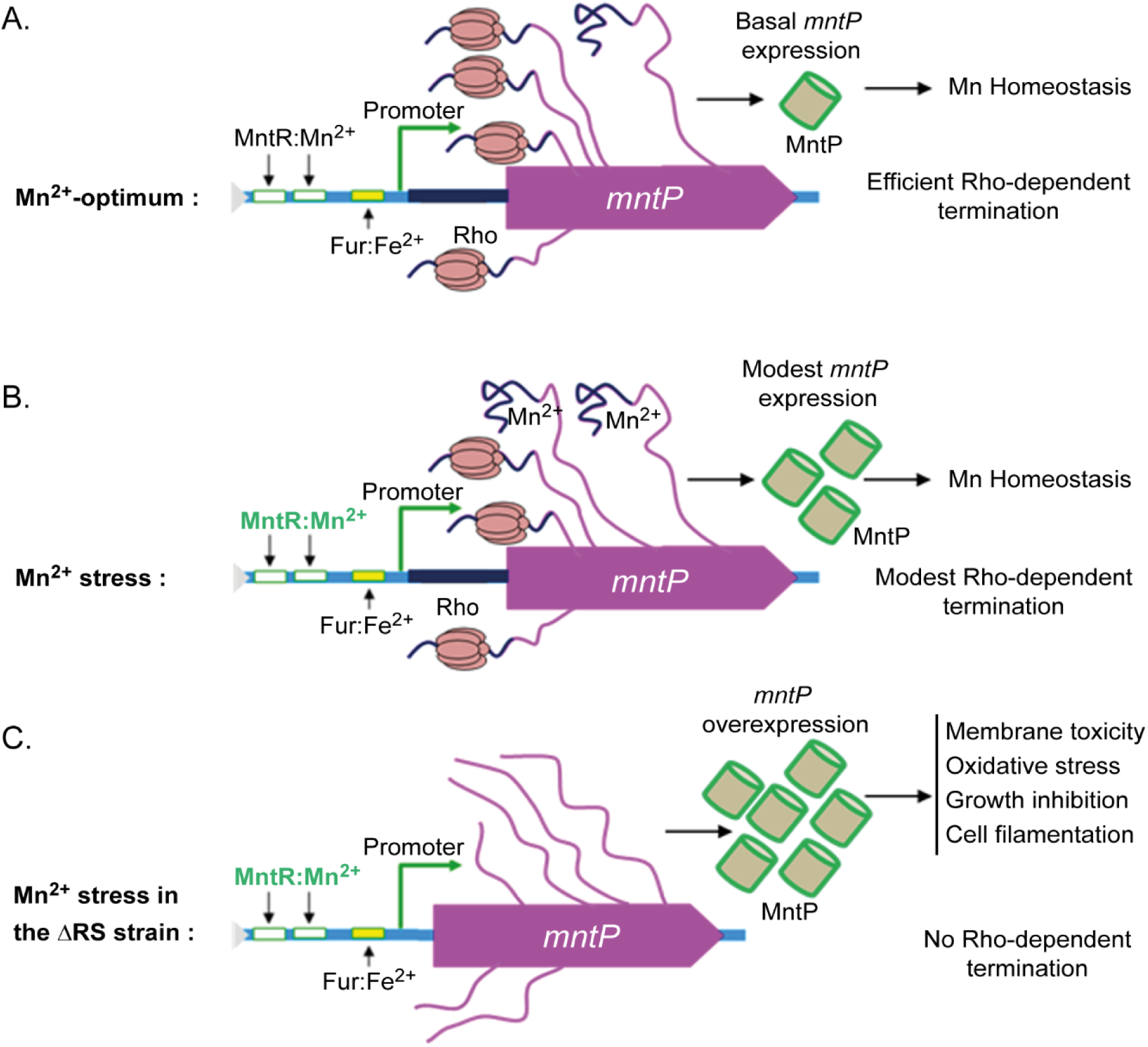
Schematic showing the regulation of *mntP* expression by Rho and Mn. **A.** At optimum cellular Mn, Rho prematurely terminates transcripts at the *mntP* locus. Besides, MntR and riboswitch-mediated regulations also remain silent. All these aspects ensure a basal level of *mntP* expression for Mn homeostasis. **B.** Under Mn stress, the active MntR: Mn^2+^ regulator, and the activated riboswitch, upregulate MntP to some extent. However, the majority of transcripts are still terminated by Rho prematurely, causing modest level of MntP expression. **C.** In case of the ΔRS strain, the absence of riboswitch RNA evades the Rho-dependent termination. Mn stress further activated MntR regulator to upregulate MntP. Therefore, MntP is overexpressed uncontrollably, causing membrane protein toxicity.

The mechanism of cytoplasmic membrane protein toxicity is multifold. Membrane protein overexpression may exhaust the limited membrane space, leaving little room for others to be integrated, or may affect enzyme functions, especially those involved in oxidative phosphorylation/ATP synthesis ^39,40,42^. Besides, such overexpression may inhibit some crucial cellular functions by nonspecific electrostatic interaction with other membrane proteins ^53^. From our study, it appears that MntP overexpression perturbed membrane biology causing elevated ROS production during oxidative phosphorylation. The ROS production might further render slow growth and cell filamentation phenotypes, as observed.

## Materials and Methods

### Bacterial strains, growth conditions, constructs, and plasmids

The wild-type BW25113 (WT) and the knockout mutant strains of *E. coli* from the KEIO collection used in this study are listed (Table 1). The mutant alleles were freshly introduced into the WT genome by P1 phage transduction. The FLP-recombination and P1 phage transduction methods were employed to generate double and triple mutants ^38^. The mutant alleles were subjected to PCR amplification and Sanger sequencing to authenticate the genotypes. Δ*RS*::*kan^R^* and Δ*RS* strains were constructed by deleting riboswitch region of *mntP* using pKD4 plasmid, RiboS1/S3 oligonucleotides and λ-red recombination system ^38^.

**Table 1.** The list of strains and plasmids used in this work.

| Strains and plasmids | Genotype/Features | References |
| --- | --- | --- |
| **Strains** |  |  |
| BW25113 | *Escherichia coli*; *rrnB3ΔlacZ4787hsdR514Δ(araBAD)*<br>567 *Δ(rhaBAD)*568 *rph-1* | <sup>55</sup> |
| *ΔmntP* | BW25113, *ΔmntP::kan<sup>R</sup>* | <sup>55</sup> |
| *ΔmntR* | BW25113, *ΔmntR::kan<sup>R</sup>* | <sup>55</sup> |
| *ΔmntH* | BW25113, *ΔmntH::kan<sup>R</sup>* | <sup>55</sup> |
| *ΔRS::kan<sup>R</sup>* | BW25113, *ΔRS::kan<sup>R</sup>*; (*mntP* riboswitch replaced with *kan<sup>R</sup>*) | This study |
| *ΔRS* | BW25113, *ΔRS::scar*; (*kan<sup>R</sup>* was removed by FLP recombinase) | This study |
| *ΔmntR ΔRS* | BW25113, *ΔmntR::kan<sup>R</sup>*, *ΔRS::scar* | This study |
| *ΔmntR ΔmntH ΔRS* | BW25113, *ΔmntR::kan<sup>R</sup>*, *ΔmntH::kan<sup>R</sup>*, *ΔRS::scar* | This study |
| BL21 (DE3) | F<sup>-</sup> *ompT hsdS<sub>B</sub> (r<sub>B</sub><sup>-</sup>, m<sub>B</sub><sup>-</sup>) gal dcm* (DE3) | Lab collection |
| C43 (DE3) | A BL21 (DE3) derivative | A gift from Dr. K. G. Thakur <sup>44</sup> |
| Lemo (DE3) | BL21(DE3) containing the Lemo System | A gift from Dr. K. G. Thakur <sup>45</sup> |
| **Plasmids** | **Features** | **References** |
| pET28a (+) | *kan<sup>R</sup>*; T7-promoter; IPTG inducible | Novagen |
| pET-*mntP* | *mntP* without a stop codon cloned in *NcoI* (blunted) and *HindIII* sites of pET28a (+) vector | This study |
| pAH125 | *kan<sup>R</sup>*; LacZ reporter | A gift from Prof. R. Chaba <sup>54</sup> |
| pVenus | *kan<sup>R</sup>*; YFP reporter | <sup>54</sup> |
| pINT | *amp<sup>R</sup>*; λ-integrase | A gift from Prof. R. Chaba <sup>54</sup> |
| pKD4 | *amp<sup>R</sup>*; *kan<sup>R</sup>* cassette is flanked by FLP recombinase | <sup>38</sup> |
| pT7A1-*trp t'* | T7A1-*trp t'* cassette was cloned in pGEM®-T vector (Promega) | This study |
| pBAD/*Myc*-His A | *amp<sup>R</sup>*; cloning vector | Invitrogen |
| pBAD-*mntP* | *mntP* cloned in *NcoI* (blunted) and *HindIII* sites of pBAD/*Myc*-His A vector | This study |
| NOTE: *kan<sup>R</sup>*, kanamycin resistance; *amp<sup>R</sup>*, ampicillin resistance. |  |  |

The DNA cassettes for the construct 1, 2, 3 and 4 strains were made by fusion PCRs using the appropriate pairs of oligonucleotides from Supplementary table 1. The DNA cassettes were cloned into pAH125 or pVenus vectors to get the desired transcriptional fusions ^54^. The cloned plasmids were then integrated into the genome of the WT strain using pINT helper plasmid and screened for single integrants, as described ^54^.

The plasmid pBAD/*Myc*-His A (pBAD) (Invitrogen) and pET28a (+) (Novagen) were used for *mntP* cloning to generate pBAD-*mntP* and pET-*mntP* expression vectors. PCR products were generated using GK1/GK2 or GK1/GK3 primer pairs and digested with the *HindIII* enzyme. The vectors were digested with *NcoI* enzyme, blunted with Klenow enzyme, and further digested with *HindIII* enzyme. Digested vectors and PCR products were ligated and transformed to generate pBAD-*mntP* and pET-*mntP* expression vectors.

### Growth Conditions

The bacterial growths were performed in buffered LB broth or media (pH 7.0) at 37°C. Growth curve analyses were done using a Bioscreen C growth analyzer (Oy growth curves Ab Ltd), as described ^9^. For other assays, the overnight culture of *E. coli* WT cells was diluted 100-fold in the fresh LB medium with or without supplemented 8 mM Mn unless otherwise specified. The cultures were grown for 1.5 hours at 37 °C till the O.D._600_ reached about 0.3. BCM (100 µg/ml) was added at that point and allowed to grow further for 2.5 hours at 37°C. The bacterial pellets were collected, and different assays were performed.

### Serial dilution and Spot assays

Overnight cultures of the *E. coli* mutants were serially diluted, and 5 µl were spotted on LB-agar plates supplemented with or without 1 mM and 2 mM Mn to visualize the relative sensitivity while growing at 37°C for 12 hours. 2 mM Mn nicely resolved the growth differences between the strains (Figure 4A).

### qPCR analyses

The bacterial cell pellets were collected and quickly washed with 1X PBS. Total RNAs were isolated by TRIzol reagent and a bacterial RNA isolation Kit (Qiagen). DNase I treatment was done. The integrity and quality of the RNA was visualized in a 1% agarose gel. The RNA concentration was determined by a UV-1800 Shimandzu UV-spectrophotometer. 200 ng of RNA samples, primer pairs (Supplementary table 1), and GoTaq 1-Step RT-qPCR System (Promega) were used for qPCR assays. At least three independent assays were conducted. The change in fold expression in the treated samples was calculated by the ΔΔC_T_ method using *betB* mRNA qPCR as a control.

### in vitro transcription assay

The in vitro transcription using biotinylated T1 and T2 templates was performed, as described previously (21, 25). Briefly, 20 nM RNAP, 40 nM templates (T1 or T2), 10 µM purified GTP and ATP, 25 µCi of [α-^32^P] CTP (BRIT, India), and 10 µM trinucleotide ApUpC (Oligos Etc.) were dissolved in 1X transcription buffer, TB (40 mM Tris-HCl, pH 7.9, 10 mM MgCl_2_, 50 mM KCl). The reaction mixture was incubated for 5 minutes at 37°C to synthesize the initial 13 nucleotide long EC (EC_13_). 200 µM of nucleotide mixtures (of ATP, GTP, CTP, and UTP) were added to EC_13_ to synthesize RO products incubating for 10 minutes at 37°C. In the Rho-dependent termination assay, 100 µM Rho was added after synthesizing EC_13_ before adding the nucleotide mixtures. Whenever required, the slurry of streptavidin beads equilibrated with 1X TB was mixed with EC_13_ and incubated for 5 minutes. Unbound substrates were removed by washing with 1X TB, and reactions were continued with or without Rho. After completion of the reaction, the supernatant portion was separated, and streptavidin beads/pellets was suspended in an equal volume of buffer. Both supernatant and pellet fractions were mixed with formamide loading dye and run on a 6% urea-PAGE. The gels were exposed to the phosphor imager screen, and Typhoon Scanner was used to capture the images.

### **β**-galactosidase assay

For β-galactosidase assays, the bacterial strains (constructs 1, 3 and 4) were grown in the presence or absence of BCM, and Mn, as described above. The cell pellets were collected and gently washed twice with Z-buffer (60 mM Na_2_HPO_4_, 40 mM NaH_2_PO_4_, 10 mM KCl, and 1 mM MgSO_4_), and the β-galactosidase assays were performed, as described ^9^.

### YFP reporter fluorescence assay

The overnight cultures were diluted 100 folds in LB broth and grown in the presence or absence of BCM and Mn, as stated above. The cell pellets were collected and washed with 1X phosphate buffer saline (PBS), and then resuspended in 1X PBS. The flow cytometry experiments were performed using a BD FACS accuri instrument for 0.1 million cells using an FL1 laser. The MFI values of three independent experiments with s.d. were plotted.

### Intracellular ROS detection

The overnight primary cultures were diluted 100 folds in LB broth and grown in the presence or absence of 2 mM Mn, and 100 µg/ml BCM, as stated above. Cell pellet was washed with 1X PBS and divided into three different fractions having approximately equal number of cells. Of the three, one fraction was stained by 2 µM DHE, the other by 10 µM H_2_DCFDA for one hour, the third fraction was dissolved in 1X PBS. The data was acquired using FACS accuri (BD) at FL1 laser for H_2_DCFDA and FL2 laser for DHE for 0.05 million cells. Mean fluorescence intensity (MFI) values obtained from three different experiments were plotted after deducting the background fluorescence values.

### Confocal microscopy analyses

The cell morphologies and filamentations were visualized using an in-house confocal Nikon confocal microscope. The *E. coli* mutants were grown, as described above in the presence or absence of Mn. The cell pellets were collected and washed with PBS. The cells were fixed with 4% formaldehyde and examined. The images were captured, and the relative cell lengths (50 cells for each condition) were measured by Image J. software. The mean and s.d. values were calculated.

### ICP MS analyses

The *E. coli* WT strain was grown in the presence or absence of 100µg/ml of BCM. The cell pellets were collected, and ICP-MS analyses were done, as described ^9^.

## Acknowledgments

The authors are grateful to Dr. Rachna Chaba lab, IISER Mohali, India, and Dr. K. G. Thakur, CSIR IMTECH, India for generously providing some plasmid and strains listed in Table 1. A.P., A.K. and K. are DBT fellow, DST-Inspire fellow and CSIR SRF fellow, respectively. R.K.M. and D.G. are UGC fellow and CSIR fellow, respectively. BP is Senior Research Scientist, NYUMC. EN is Julie Wilson Anderson Professor, and HHMI fellow. This work received financial support from intramural grants of CSIR (MLP-0042) and CSIR-IMTECH (OLP-0190), and by an extramural grant to D.D. from SERB (CRG/2019/001174), Department of Science and Technology (DST).

## Author contributions

A.P. and A.K. made constructs, *lacZ* reporters with riboswitch. K performed qPCR analyses, made construct with *trp t’* terminators. A.P, A.K. and K performed most of the experiments including reporter assays. R.K.M. conducted YFP reporter assays and helped all three first authors. A.P. and D.G. purified the proteins and performed in vitro transcription assays. G.K. made *mntP* clones and mutants, performed growth analyses and western blotting experiments. B.P. and E.N. analyzed and substantially modified the manuscript. E.N. and D.D. analyzed and interpreted the data. E.N. approved the submitted version. D.D. designed the experiments and supervised A.P., A.K., K., R.K.M., D.G., and G.K.

## Supplementary materials

**Supplementary Figure 1.**
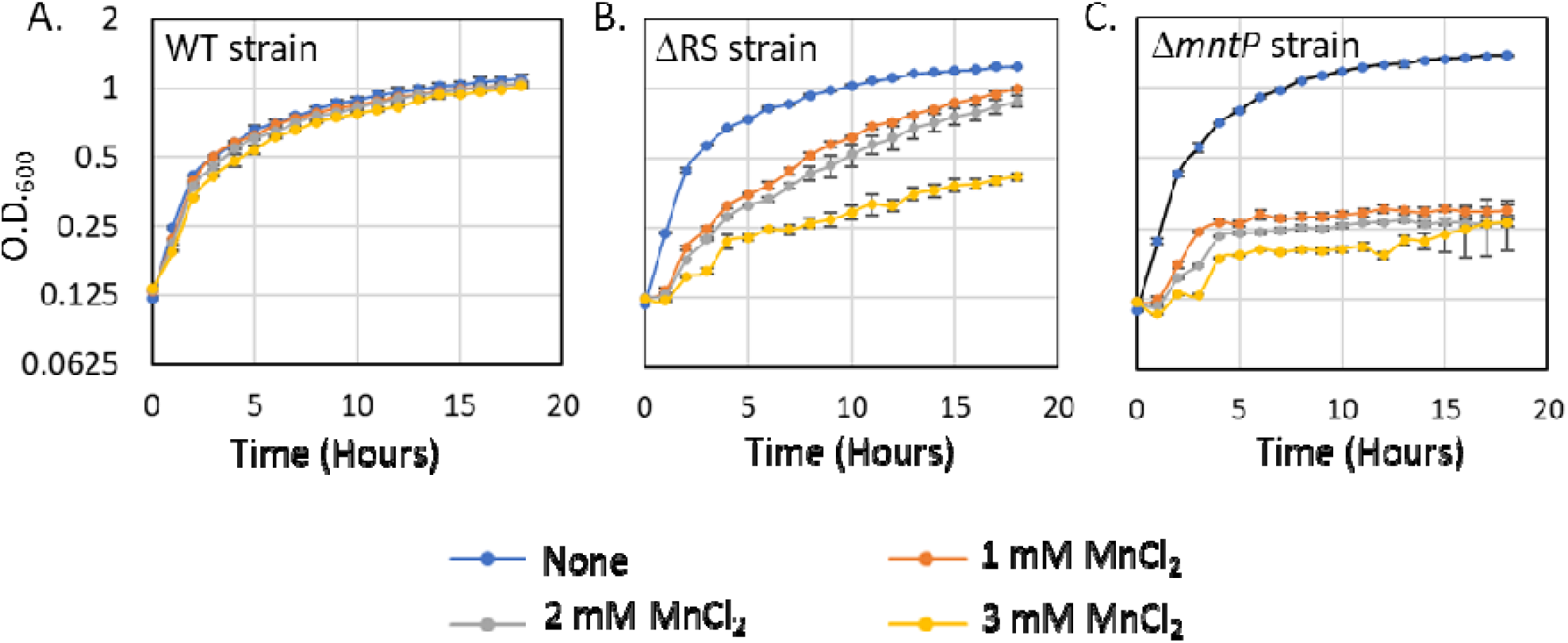
The WT (**A**), ΔRS (**B**), Δ*mntP* (**C**) strains were grown in absence or the presence of different concentrations of manganese to compare the manganese sensitivity. Roughly, 3 mM manganese-fed ΔRS strain and 1 mM manganese-fed Δ*mntP* strain exhibit similar level of growth retardation.

**Supplementary Figure 2.**
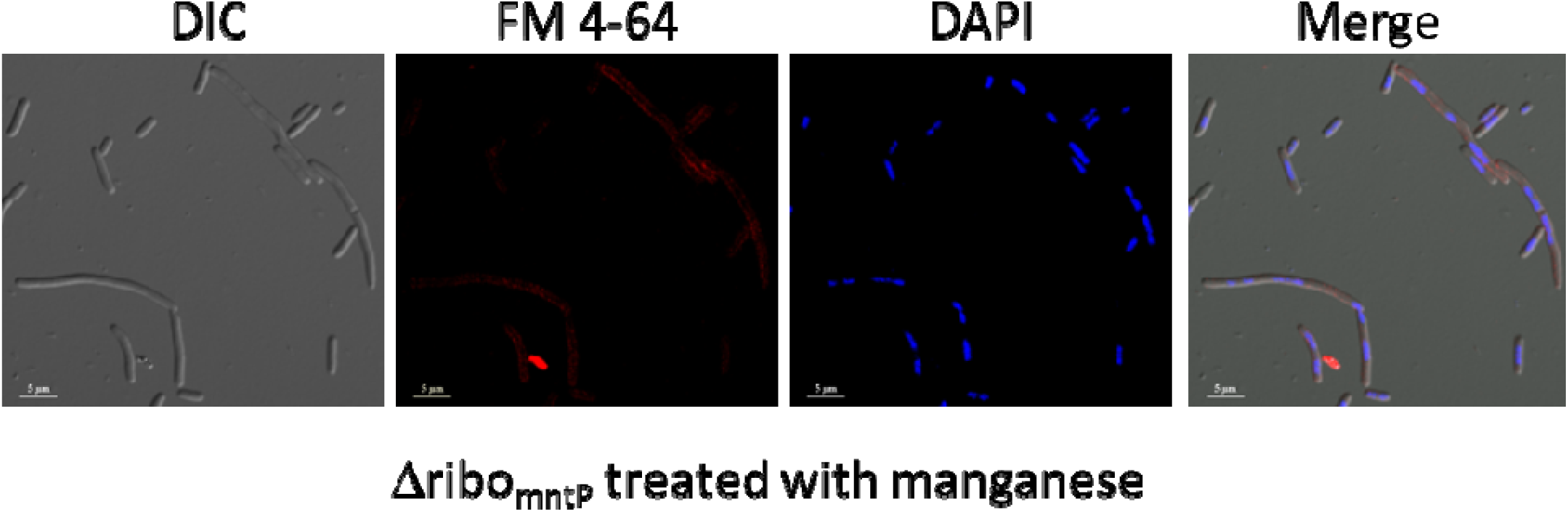
Mn-treated Δ*RS* cells were stained with FM-4-64 and DAPI to show that the filamentous cells were without any septum and discrete chromosomal DNA units were arranged throughout the filamented cells.

**Supplementary Figure 3.**
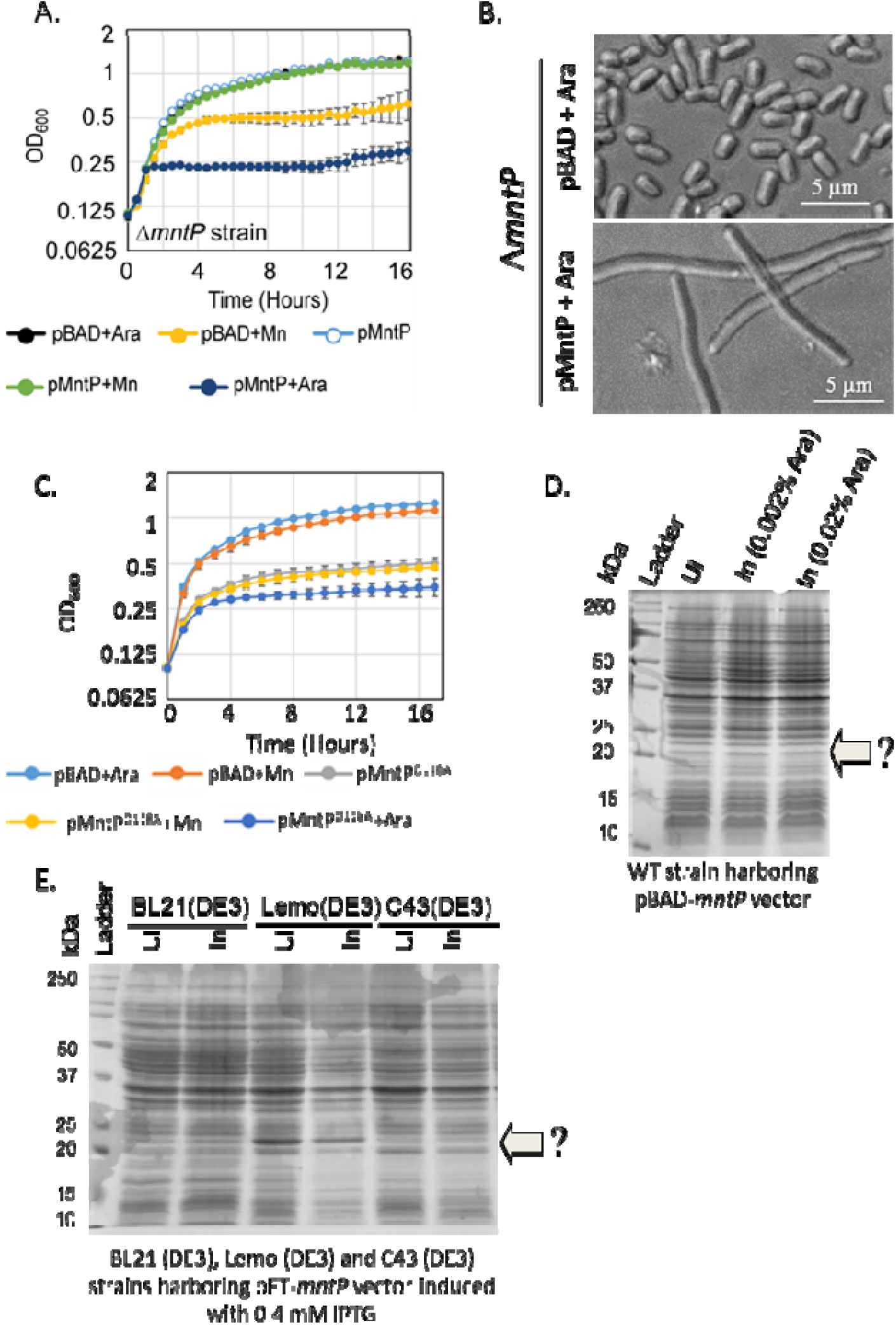
A. The multicopy induction of MntP expression by arabinose (0.002%) is highly toxic to the Δ*mntP*, a manganese sensitive strain. The overexpression related toxicity is higher than the manganese toxicity at 1 mM, as observed. The calculated mean ± s.d. values from three independent experiments were plotted. **B**. The multicopy induction of MntP also leads to cell filamentation. **C.** The multicopy induction of MntP^D118A^, which is also manganese sensitive, by arabinose (0.002%) is highly toxic to the Δ*mntP* cells. The calculated mean ± s.d. values from three independent experiments were plotted. **D.** *mntP* was induced with two different concentration araboinose from WT strain harboring pBAD-*mntP* vector, and SDS-PAGE was performed. No overexpressed band against MntP was detected after coomassie staining of the gel. The arrow with the question mark denotes molecular weight position (∼20 kD) where MntP would be visualized if overexpressed above detection level. **D**. His-tagged *mntP* was induced with IPTG from the different *E. coli* strains harboring pET-*mntP* vector, and SDS-PAGE was performed. The overexpressed band against MntP-6X-his remained undetected after coomassie staining of the gel. The arrow with the question mark denotes molecular weight position (∼22 kD) where MntP-6X-his would be visualized if overexpressed enough.

**Supplementary Table 1.**
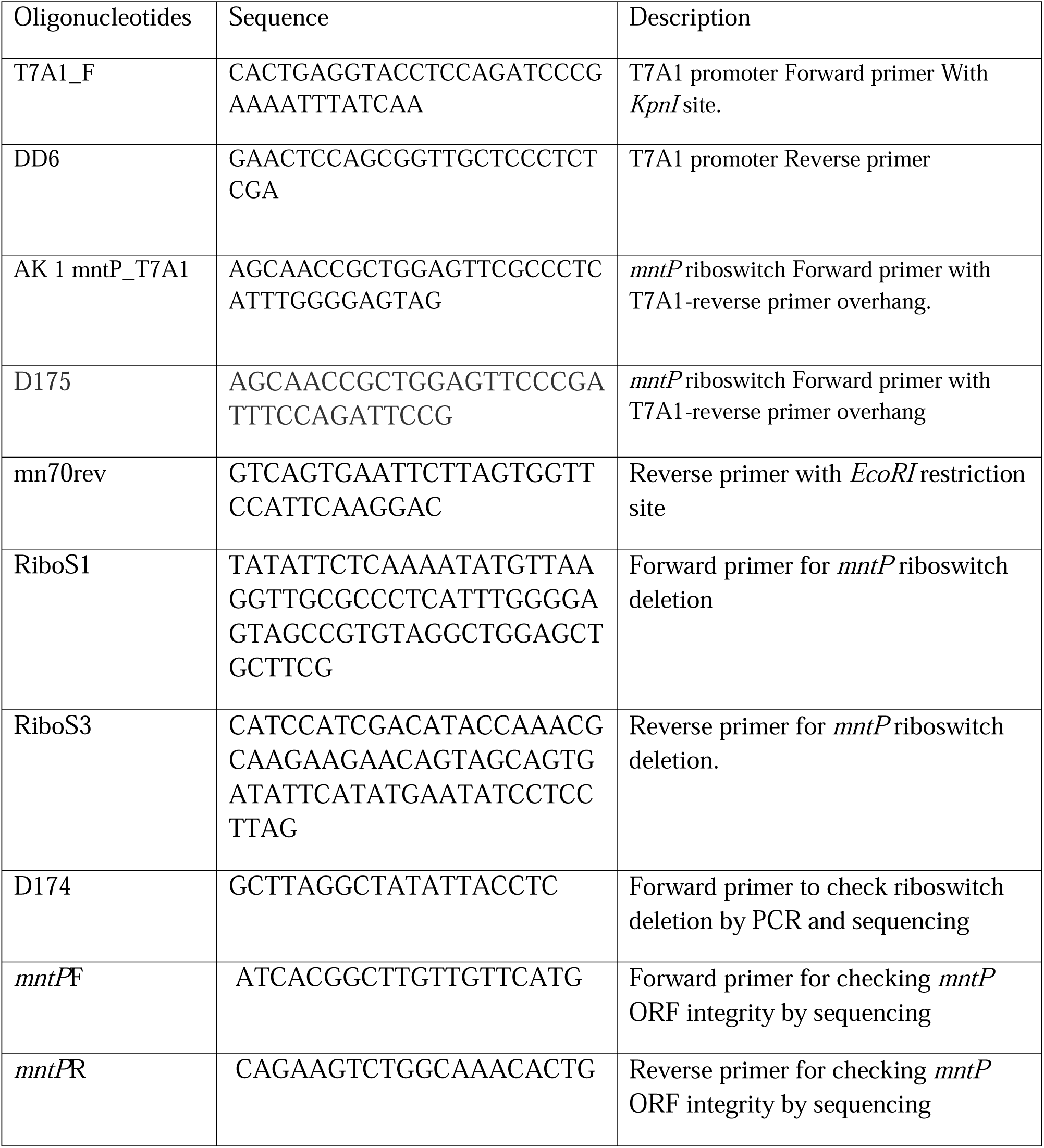
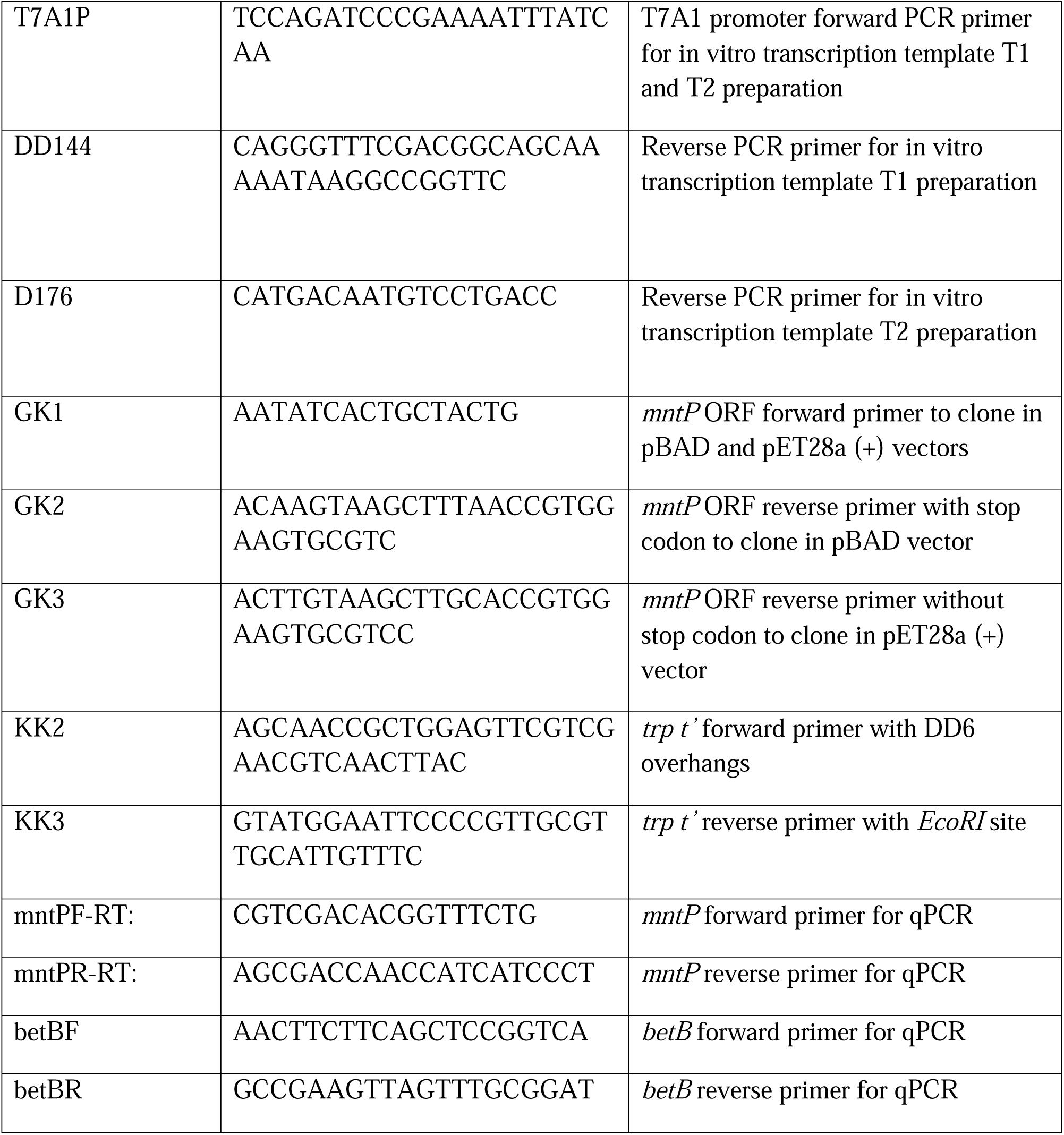
List of oligonucleotides used in this study.

